# Massively parallel mutant selection identifies genetic determinants of *Pseudomonas aeruginosa* colonization of *Drosophila melanogaster*

**DOI:** 10.1101/2023.11.20.567573

**Authors:** Jessica Miles, Gabriel L. Lozano, Jeyaprakash Rajendhran, Eric V. Stabb, Jo Handelsman, Nichole A. Broderick

## Abstract

*Pseudomonas aeruginosa* is recognized for its ability to colonize diverse habitats and cause disease in a variety of hosts, including plants, invertebrates, and mammals. Understanding how this bacterium is able to occupy wide-ranging niches is important for deciphering its ecology. We used transposon sequencing (Tn-Seq, also known as INSeq) to identify genes in *P. aeruginosa* that contribute to fitness during colonization of *Drosophila melanogaster*. Our results reveal a suite of critical factors, including those that contribute to polysaccharide production, DNA repair, metabolism, and respiration. Comparison of candidate genes with fitness determinants discovered in previous studies of *P. aeruginosa* identified several genes required for colonization and virulence determinants that are conserved across hosts and tissues. This analysis provides evidence for both the conservation of function of several genes across systems, as well as host-specific functions. These findings, which represent the first use of transposon sequencing of a gut pathogen in *Drosophila*, demonstrate the power of Tn-Seq in the fly model system and advance existing knowledge of intestinal pathogenesis by *D. melanogaster,* revealing bacterial colonization determinants that contribute to a comprehensive portrait of *P*. *aeruginosa* lifestyles across habitats.

**Importance:** *Drosophila melanogaster* is a powerful model for understanding host-pathogen interactions. Research with this system has yielded notable insights into mechanisms of host immunity and defense, many of which emerged from analysis of bacterial mutants defective for well-characterized virulence factors. These foundational studies – and advances in high-throughput sequencing of transposon mutants – support unbiased screens of bacterial mutants in the fly. To investigate mechanisms of host-pathogen interplay and exploit the tractability of this model host, we used a high-throughput, genome-wide mutant analysis to find genes that enable a pathogen, *P. aeruginosa*, to colonize the fly. Our analysis reveals critical mediators of *P. aeruginosa* establishment in its host, some of which are required across fly and mouse systems. These findings demonstrate the utility of massively parallel mutant analysis and provide a platform for aligning the fly toolkit with comprehensive bacterial genomics.

## Introduction

*Drosophila melanogaster* has long been an effective model organism for investigating bacterial infection and host-microbe interactions. Traditionally, these studies emphasized host responses to pathogens delivered by septic injury, but the more recent identification of microbes that infect flies following ingestion has facilitated the study of enteric pathogens (1,2). Studies of entomopathogens, such as *Pseudomonas entomophila* and *Pectobacterium carotovora*, and broad host-range pathogens, such as *Serratia marcescens* and *Pseudomonas aeruginosa*, have elucidated global mechanisms of gut homeostasis and host defense (3–12). Although comprehensive mutant analyses of *Francisella novicida* and *Mycobacterium marinum* demonstrate the potential of *D. melanogaster* as a host for forward genetic screens of bacterial pathogens, a genome-wide screen of an ingested pathogen has not been reported (13,14). Recent advances in signature-tagged mutagenesis offer additional methods for a comprehensive genetic dissection of the fitness determinants that enable ingested bacteria to survive within the fly.

One of these techniques, transposon sequencing (Tn-Seq) has emerged as a powerful tool for identifying genes that enable bacteria to colonize a variety of vertebrate and invertebrate hosts (15–18). This approach combines traditional transposon-mutant analysis with next-generation sequencing in a single-selection, high-throughput screen. Tn-Seq can be used to evaluate changes in the frequency of a mutated gene within a population, providing quantitative data that can be evaluated statistically. As a result, this technique can identify mutants that have either increased or decreased fitness when presented to the host in a pool. Tn-Seq also imparts population-level data on the relative fitness among mutants, which is useful for monitoring subtle phenotypes.

Here, we used Tn-Seq to identify genes in *P*. *aeruginosa* that contribute to colonization of the fly during oral infection. We constructed a saturated library of *P. aeruginosa* mutants, administered mutant pools of optimal complexity to flies, and used massively parallel sequencing to reveal mutants that were negatively selected during administration of the library and after consumption by the fly. Some of the putative colonization factors were identified and characterized previously as virulence factors in other infection models, thereby validating this genetic approach. Other putative colonization factors revealed by our screen include genes involved in DNA repair, metabolism, and nutrition. Notably, many determinants of bacterial fitness in the fly are homologs of genes required for *P. aeruginosa* viability in other systems. In sum, these findings deepen our understanding of *P*. *aeruginosa* infection, underscore the utility of the fly model, and validate the use of Tn-Seq in *D*. *melanogaster*.

## Results

### Generating an input library for Insertion Sequencing (Tn-Seq)

We generated a transposon-mutant library of *P. aeruginosa* containing over 47,000 independent insertions using a mariner-based transposon (18–20), which were well distributed across the genome (Figure 1A). After excluding insertions in the distal 10% of open reading frames, we identified 520 genes with no insertions (Figure 1B; 21), a set with a high concordance among technical replicates of the library. This transposon library constituted the “input population” for the following experiments in flies.

**Figure 1:**
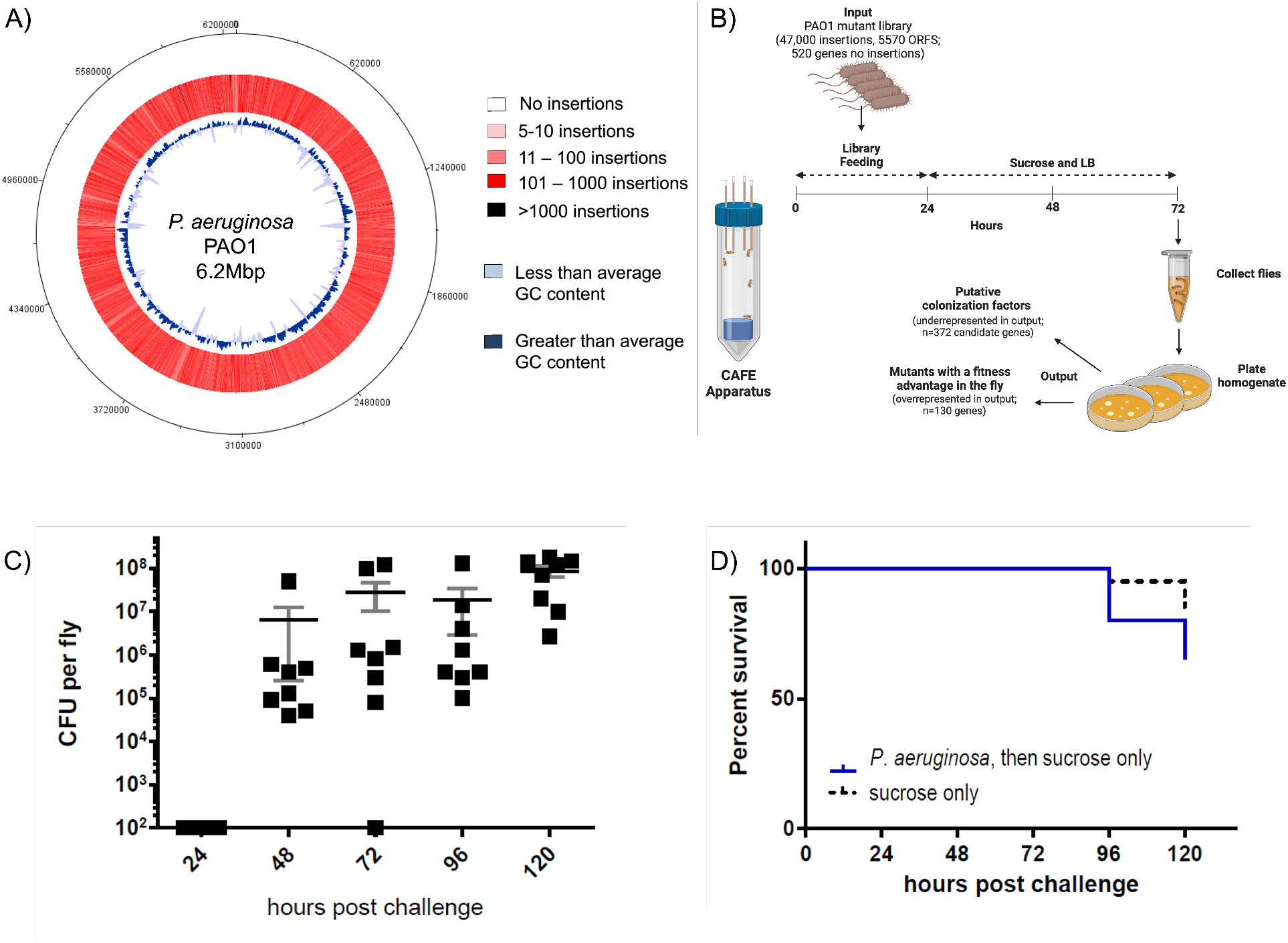
Using the Tn-Seq platform to identify colonization factors of *P. aeruginosa* in the fly. **(A)** A highly saturated *P. aeruginosa* transposon library was constructed, prepared for Tn-Seq, and characterized (“input”). Approximately 62,000 insertions – 47,000 of them unique (37,000 in ORFs and 10,000 in intergenic regions) – were mapped to the *P*. *aeruginosa* PAO1 genome. Color intensity increases with increasing number of insertion (white < 10 insertions; pink = 11-100 insertions; red =101-1000 insertions; sites with >1000 insertions are black). GC content is also shown (dark blue: greater than average; light blue: less than average; GC content = 66.56%). **(B)** 3-7-day-old female Canton-S flies housed in capillary feeders (CAFEs; a cartoon is shown above) were fed a population of PAO1 transposon mutants (input) for 24 hours, then were switched to a sucrose-LB broth suspension for an additional 48 hours (n=250 flies). After the input was administered to flies, fly homogenate was cultured and *P. aeruginosa* mutants were recovered, prepared for Tn-Seq, and characterized (“output”). Genes underrepresented after passage through the fly were considered “putative colonization factors.” Genes with no insertions in the input were not analyzed in the output. Flies were sampled 72 hours post-challenge. Six independent replicates were performed. **(C and D)** 3-7-day-old female were fed on a 10^5^ CFU/mL suspension in 5% sucrose for 24 hours. Then, they were transferred to 5% sucrose in LB and remained on the sucrose solution for the duration of the experiment. Flies were sampled daily for the 5-day period. For CFU per fly **(C)**, means and SEM are shown. Limit of detection is at axis (100 CFU per fly). n= 8 flies per group per day sampled. For fly survival **(D)**, n = 20 flies.

### Screening a transposon library in *Drosophila melanogaster* using a capillary feeder

We fed the input library to flies *ad libitum*, utilizing a capillary feeder (Figure 1B, 22), which facilitated monitoring the volume of the library suspension ingested by flies. We assessed *P. aeruginosa* population bottlenecks in pools of wild-type and mutant strains mixed at different ratios to confirm that we could reproducibly recover mutants fed to flies even if they are a small portion of the input. Our goal was to use *P. aeruginosa* mutant pools that enabled establishment of a large number of mutants in the fly (∼10^3^) without severe stochastic loss from colonization bottlenecks. We determined that an inoculum of 10^5^ CFU/mL fed to and recovered from 250 flies represented a sufficiently large and complex input pool to screen mutants in the library without significant bottlenecks between the input and cells recovered from the host. We also determined that the timing of our feeding and recovery minimized infection lethality while maximizing the size of the population of *P. aeruginosa* at the time of collection (Figure1C and D). Six replicates of 250 flies each, were given access to the library for 24 hours followed by access to sterile sucrose solution for an additional 48 hours. For each replicate, *P. aerouginosa* mutants established in fly guts (“output population”) were recovered from surface-sterilized homogenized flies by culturing on LB agar (Figure 1B) prior to DNA extraction and Tn-Seq analysis.

### Identification of genes in *P. aeruginosa* that contribute to fitness during colonization of the fly

We identified 372 candidate genes that contribute to fitness during colonization of the fly (“*in vivo*”) (Table S2 in 21). To distinguish genes that are important for colonization of flies (“*in vivo*”) from genes that are critical for bacterial survival across the feeding portion of the experiment (“*in vitro”*), we also characterized an input population that was maintained in capillary feeders, but not exposed to flies (Table S3 in 21). We identified 379 candidate genes that contribute to fitness in this condition (Table S3 in 21). In comparing these “*in vivo*” and “*in vitro*” datasets, we found 294 genes depleted in both conditions, but also identified candidates unique to each condition, indicating that there are selection pressures that differ between feeding in the capillary and in the fly (Figure 2A, Tables S4 and S5 in 21).

**Figure 2.**
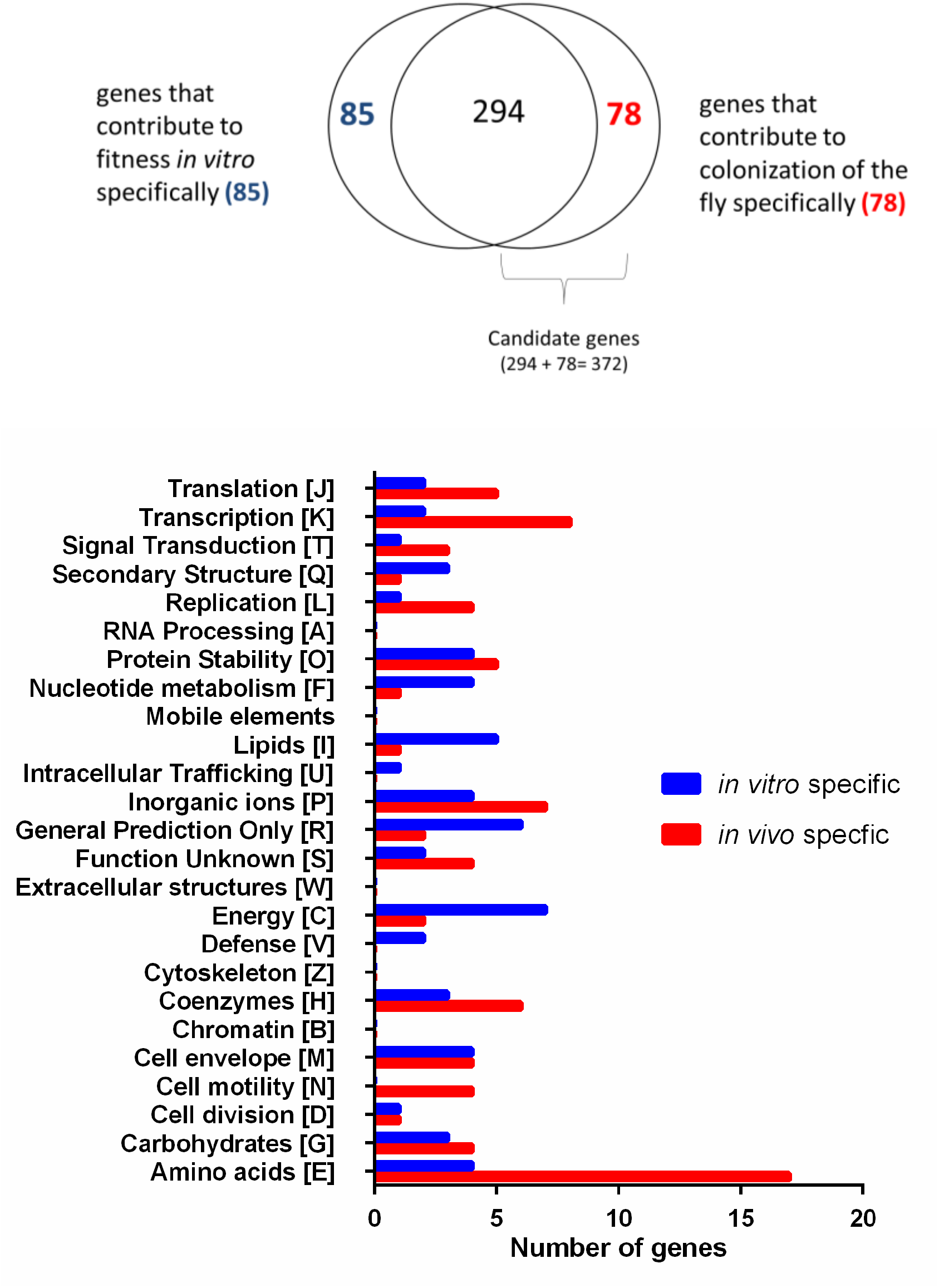
Comparison of *P*. *aeruginosa* fitness determinants *in vitro* and *in vivo*. **(A)** After characterizing a mock output population that was subjected to experimental conditions, but not administered to flies, we compared genes depleted in that group (n=379) to the output population recovered from flies (n=372). Eighty-five genes were depleted only *in vitro*; 78 genes were depleted only *in vivo*. Genes that were negatively selected *in vivo*, but not *in vitro* were termed putative “colonization-specific factors.” Nearly 300 genes were negatively selected both while the library was administered to flies (*in vitro*) and during passage through the fly (*in vivo*). **(B)** Genes depleted specifically during administration of the library tended to be categorized as contributing to secondary structure (COG category Q), nucleotide metabolism and transport (F), lipid metabolism (I), and energy production and conversion (C), whereas genes depleted specifically in the fly were tended to be categorized as contributing to transcription, translation, signal transduction, replication and repair, cell motility, and amino acid metabolism and transport. COG categories B (chromatin structure and dynamics), W (extracellular structures), Z (cytoskeleton), and mobile elements were not represented among these genes specifically depleted in either condition. COG = Clusters of Orthologous Groups. Full lists of genes can be found in Tables S2-S5 in reference 21.

After categorizing these genes based on Clusters of Orthologous Groups (COG) designations, we discovered that categories representing synthesis of secondary metabolites, nucleotide metabolism and transport, lipid metabolism, and energy production and conversion were underrepresented in the *in vitro* output, whereas genes categorized as contributing to transcription, translation, signal transduction, replication and repair, cell motility, and amino acid metabolism and transport were underrepresented in the *in vivo* output (Figure 2B). However, a number of fitness determinants were shared between the two conditions (Table S6 in 21); based on our observations, this overlap encompasses global requirements for viability (Figure 2A). We interpreted this overlap as suggesting that the feeding apparatus and selection in fly (on sterile sucrose for 48 hours after the 24-hour library feeding period) were likely equally restricted nutritionally and thus these genes are important for colonization in such conditions. As the presence of the host did not expand representation of these *in vitro* depleted genes, we included them in our downstream analysis as putative colonization genes. The annotations of genes that contribute to fitness during colonization of the fly revealed functions that were shared among candidates, including virulence; synthesis of flagella and surface polysaccharides; DNA repair; synthesis of nucleotides, amino acids, and cofactors; and aerobic respiration (Table 1).

**Table 1.** Annotated functions of colonization factors identified through Tn-Seq screening. Based on annotation, candidate genes contribute to virulence, production of alginate and psl polysaccharides, DNA repair, biosynthesis of small molecules (nucleotides, amino acids, and B vitamins), and respiration, among other functions.

| Role (annotation) | Gene ID | Gene | COG Category | Virulence Factor? | Fly specific |
| --- | --- | --- | --- | --- | --- |
| Virulence Factors | PA0336 | *ygdP* | Defense mechanisms [V] | reported (55) |  |
|  | PA4229 | *pchC* | Secondary structures [Q] | VFDB |  |
|  | PA1776 | *sigX* | Transcription [K] |  |  |
|  | PA1777 | *oprF* | Cell envelope [M] | reported (25) |  |
|  | PA2397 | *pvdE* | Inorganic ions[P] | VFDB |  |
|  | PA4494 | *roxS* | Signal transduction [T] | reported (28) |  |
|  | PA4856 | *retS* | Signal transduction [T] | reported (56) |  |
| Flagella | PA1094 | *fliD* | Cell motility [N] | VFDB | + |
|  | PA1077 | *flgB* | Cell motility [N] | VFDB | + |
|  | PA1078 | *flgC* | Cell motility [N] | VFDB | + |
|  | PA1453 | *flhF* | Cell motility [N] | VFDB |  |
|  | PA1454 | *fleN* | Cell motility [N]; Cell division [D] | VFDB |  |
|  | PA1455 | *fliA* | Transcription [N] | VFDB | + |
|  | PA1456 | *cheY* | Signal transduction [T] |  |  |
|  | PA1461 | *motD* | Cell motility [N] | VFDB | + |
|  | PA3351 | *flgM* | Transcription [K] | VFDB |  |
| Polysaccharides | PA5322 | *algC* | Carbohydrates [G] | VFDB |  |
|  | PA5262 | *algZ/fimS* | Signal transduction [T] | VFDB |  |
|  | PA0762 | *algU* | Transcription [K] | VFDB |  |
|  | PA0763 | *mucA\** | Signal transduction [T] | VFDB |  |
|  | PA1727 | *mucR* | Signal transduction [T] |  |  |
|  | PA3649 | *mucP* | Cell envelope [M] |  |  |
|  | PA1801 | *clpP* | Protein stability [O] | (16) |  |
|  | PA1802 | *clpX* | Protein stability [O] | (16) |  |
|  | PA2234 | *pslD* | Cell envelope [M] |  | + |
|  | PA2235 | *pslE* | Cell envelope [M] |  | + |
|  | PA2236 | *pslF* | Cell envelope [M] |  |  |
|  | PA2239 | *pslI* | Cell envelope [M] |  |  |
| DNA repair | PA0965 | *ruvC* | Replication and repair [L] |  |  |
|  | PA0966 | *ruvA* | Replication and repair [L] |  |  |
|  | PA0967 | *ruvB* | Replication and repair [L] |  |  |
|  | PA5344 | *oxyR* | Transcription [K] | reported (57) |  |
|  | PA5345 | *recG* | Replication and repair [L] |  |  |
|  | PA0407 | *gshB* | Coenzyme metabolism [H] |  | + |
|  | PA5203 | *gshA* | Coenzyme metabolism [H] |  |  |
|  | PA3625 | *surE* | Replication and repair [L] |  | + |
|  | PA3777 | *xseA* | Replication and repair [L] |  |  |
|  | PA3617 | *recA* | Replication and repair [L] |  |  |
|  | PA4283 | *recD* | Replication and repair [L] |  |  |
|  | PA4284 | *recB* | Replication and repair [L] |  |  |
|  | PA4285 | *recC* | Replication and repair [L] |  |  |
| Biosynthetic Pathways | PA0944 | *purN* | Nucleotide metabolism [F] |  | + |
|  | PA0945 | *purM* | Nucleotide metabolism [F] |  |  |
|  | PA3108 | *purF* | Nucleotide metabolism [F] |  |  |
|  | PA4854 | *purH* | Nucleotide metabolism [F] |  |  |
|  | PA4855 | *purD* | Nucleotide metabolism [F] |  |  |
|  | PA5425 | *purK* | Nucleotide metabolism [F] |  |  |
|  | PA4756 | *carB* | Nucleotide metabolism [F] |  |  |
|  | PA4758 | *carA* | Nucleotide metabolism [F] |  |  |
|  | PA3527 | *pyrC'* | Nucleotide metabolism [F] |  |  |
|  | PA0402 | *pyrB* | Nucleotide metabolism [F] |  |  |
|  | PA0430 | *metF* | Amino acids [E] |  | + |
|  | PA3107 | *metZ* | Amino acids [E] |  | + |
|  | PA3166 | *pheA* | Amino acids [E] |  |  |
|  | PA5277 | *lysA* | Amino acids [E] |  |  |
|  | PA0025 | *aroE* | Amino acids [E] |  |  |
|  | PA5038 | *aroB* | Amino acids [E] |  |  |
|  | PA5039 | *aroK* | Amino acids [E] |  |  |
|  | PA4846 | *aroQ1* | Amino acids [E] |  |  |
|  | PA3165 | *hisC2* | Amino acids [E] |  | + |
|  | PA5140 | *hisF1* | Amino acids [E] |  |  |
|  | PA5067 | *hisE* | Amino acids [E] |  |  |
|  | PA5066 | *hisI* | Amino acids [E] |  |  |
|  | PA0035 | *trpA* | Amino acids [E] |  | + |
|  | PA0036 | *trpB* | Amino acids [E] |  |  |
|  | PA0649 | *trpG* | Amino acids [E] |  | + |
|  | PA0353 | *ilvD* | Amino acids [E] |  | + |
|  | PA4695 | *ilvH* | Amino acids [E] |  | + |
|  | PA4694 | *ilvC* | Amino acids [E] |  |  |
|  | PA4696 | *ilvI* | Amino acids [E] |  |  |
|  | PA5013 | *ilvE* | Amino acids [E] |  |  |
|  | PA5204 | *argA* | Amino acids [E] |  |  |
|  | PA5323 | *argB* | Amino acids [E] |  | + |
|  | PA0662 | *argC* | Amino acids [E] |  | + |
|  | PA3525 | *argG* | Amino acids [E] |  | + |
|  | PA3537 | *argF* | Amino acids [E] |  |  |
|  | PA5263 | *argH* | Amino acids [E] |  |  |
|  | PA3118 | *leuB* | Amino acids [E] |  | + |
|  | PA3121 | *leuC* | Amino acids [E] |  |  |
|  | PA5495 | *thrB* | Amino acids [E] |  |  |
|  | PA3736 | *hom* | Amino acids [E] |  | + |
|  | PA0316 | *serA* | Amino acids [E] |  |  |
|  | PA4565 | *proB* | Amino acids [E] |  | + |
|  | PA0381 | *thiG* | Coenzyme metabolism [H] |  |  |
|  | PA5118 | *thiI* | Coenzyme metabolism [H] |  | + |
|  | PA3975 | *thiD* | Coenzyme metabolism [H] |  | + |
|  | PA0501 | *bioF* | Coenzyme metabolism [H] |  |  |
|  | PA0500 | *bioB* | Coenzyme metabolism [H] |  |  |
|  | PA0502 | *bioH* | Coenzyme metabolism [H] |  |  |
|  | PA0420 | *bioA* | Coenzyme metabolism [H] |  |  |
| Respiration | PA1553 | *ccoO1* | Energy [C] |  |  |
|  | PA1556 | *ccoO2* | Energy [C] |  |  |
|  | PA5300 | *cycB* | Energy [C] |  |  |
|  | PA5490 | *cc4* | Carbohydrates [G] |  |  |
|  | PA2637-PA2649 | *nuoABC*<br>*DEFGHI*<br>*JKLMN* | Energy [C] |  |  |

#### Virulence factors

Colonization is vital to the establishment of infection, and we predicted that some of our candidates would have previously characterized roles in pathogenesis. To survey virulence factors, we examined the Virulence Factors of Pathogenic Bacteria Database (http://www.mgc.ac.cn/cgi-bin/VFs/compvfs.cgi?Genus=Pseudomonas) and found that several putative colonization factors identified here were also listed in that database, including genes encoding a thioesterase (PchC) and an ABC-type transporter protein (PvdE) required for synthesis of the siderophores pyochelin and pyoverdine (23, 24). Other candidates, such as the outer membrane porin OprF, did not appear in this database, but have been reported elsewhere to contribute to virulence (25). The ECF sigma factor *sigX*, which lies directly upstream of *oprF* and modulates its expression, was also important for fitness (26). Insertions in genes encoding RetS, a hybrid sensor kinase/response regulator with known roles in colonization and virulence, and RoxS, one component of the RoxS/RoxR sensor histidine kinase regulator, were similarly depleted in the output (27, 28).

#### Flagella

Mutants with insertions in several components of the flagellar apparatus were depleted in the output, indicating that flagellar function is critical for fitness in the fly. These mutants mapped in genes that encode structural components of the flagellum – such as the basal-body rod modification protein, FlgD, and the capping protein, FliD (www.pseudomonas.com). Other fitness determinants were regulators of flagella production and assembly, such as sigma factor FliA, synthesis regulator FleN, anti-sigma factor FlgM, and CheY, a global regulator of flagella production and chemotaxis (www.pseudomonas.com).

#### Surface polysaccharides

Pseudomonads are well-known exopolysaccharide producers, and the role of polysaccharides in *P. aeruginosa* biofilm production has been widely studied. We identified putative colonization factors associated with these functions such as regulators of alginate (*algC, algZ/fimS, algU, clpP, clpX*) and psl polysaccharide (*pslD, pslE, pslF, pslI*), two of the three major exopolysaccharides produced by *P. aeruginosa*, as well as the regulators *mucR*, *mucP*, and *mucA* (29, www.pseudomonas.com).

#### DNA repair factors

Previous transposon-sequencing analysis of *P. aeruginosa* has demonstrated that its defense against reactive oxygen species (ROS) contributes to survival across environments (30). The ability to withstand ROS is critical for bacterial survival in the fly intestine, where ROS are produced at low levels in response to symbiotic lactobacilli and increased in response to enteric infection (31–33). We identified several factors that mediate DNA repair in our screen, an important response to oxidative stress, including the RuvABC resolvasome; the homologous recombination proteins RecA, RecBCD, and RecG; and the survival protein SurE. Mutations in *gshB*, which encodes the antioxidant glutathione synthase; *oxyR*, the gene encoding a potent regulator of oxidative stress response genes; and *xseA*, the gene encoding exonuclease VII, were also negatively selected.

#### Biosynthetic pathways

Mutants deficient for the synthesis of amino acids, cofactors, and nucleotides were underrepresented in the output. Genes required for the synthesis of aromatic and branched chain amino acids and arginine were especially prominent among this group of candidates, as were genes essential for biotin synthesis.

#### Respiration

Aerobic respiration is the predominant mechanism of energy production for *P*. *aeruginosa* (17). Mutants lacking NADH:ubiquinone oxidoreductase (complex I) and cytochrome c oxidase (complex IV) were impaired, a phenotype that has been associated with virulence in other hosts (17). Transposon insertions in genes encoding a precursor of cytochromes c4 and c5 were also depleted.

### Global regulators of *P. aeruginosa* colonization

To identify *P. aeruginosa* colonization determinants that were functionally conserved across systems, we compared homologs of putative colonization factors that we identified to those previously reported *P. aeruginosa* PA14 and PAO1 fitness determinants in the murine intestine and an acute burn model (Figure 3, [17, 34]). We observed that, although some genes are uniquely important in each system, the loss of certain functions is detrimental in all host systems: we found a shared requirement for surface polysaccharides, DNA repair factors, biosynthetic factors, and respiration genes across habitats (Figure 3A, [17, 34]). The functions of fitness determinants in the fly were largely shared with those in murine environments, whereas each mouse system had its own requirements (Figure 3A). Notably, *cheY*, a response regulator of flagellar rotation whose function is required for chemotaxis, was critical for colonization of the fly and in both mouse infection systems (Figure 3A; [17, 34, 35]). Moreover, mutations in a small number of genes enhanced fitness *in vivo* (Table S7 in 21), including those encoding isocitrate lyase AceA, which was also enriched in the mouse intestine, and the multidrug efflux pump operon MexEF-OprN whose overproduction impairs virulence in *C. elegans* infection (Figure 3B, 17, 36).

**Figure 3.**
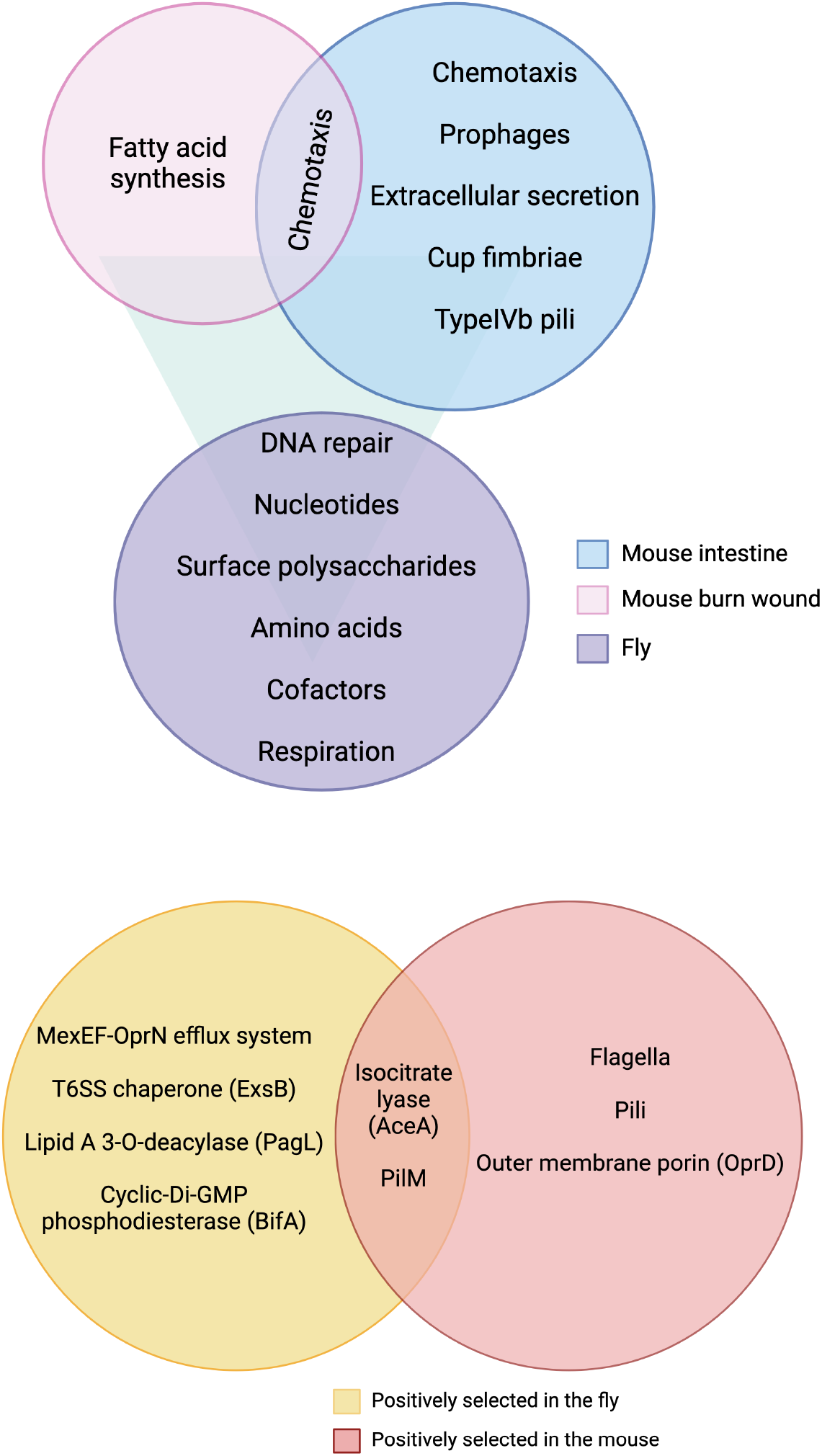
Comparison of genetic determinants of *P*. *aeruginosa* colonization in the fly and mouse from previous studies. **(A)** Alginate and psl polysaccharides, purines, pyrimidines, amino acids, cofactors, and respiration genes are critical for establishment of PAO1 and *P. aeruginosa* PA14 in invertebrate and vertebrate hosts, and at different body sites (14, 31). **(B)** Mutants in *aceA*, and *pilM* were positively selected across systems, in both PAO1 and PA14. Most of the genes encoding components of flagella and pili were positively selected in the mouse, but not the fly. Insertions in the *mexEF-oprN* operon, *exsB*, *pagL*, and *bifA*, among others, were overrepresented in the fly.

### Validation of putative colonization factors

To validate the phenotypes associated with our colonization candidates, we screened four mutants in 1:1 competition assays with wild-type *P. aeruginosa* in the fly (Table 1, Figure 4). In all cases, we found that these mutants were underrepresented relative to wild type upon recovery from host flies, thereby indicating that the Tn-Seq screen identified authentic colonization factors.

**Figure 4.**
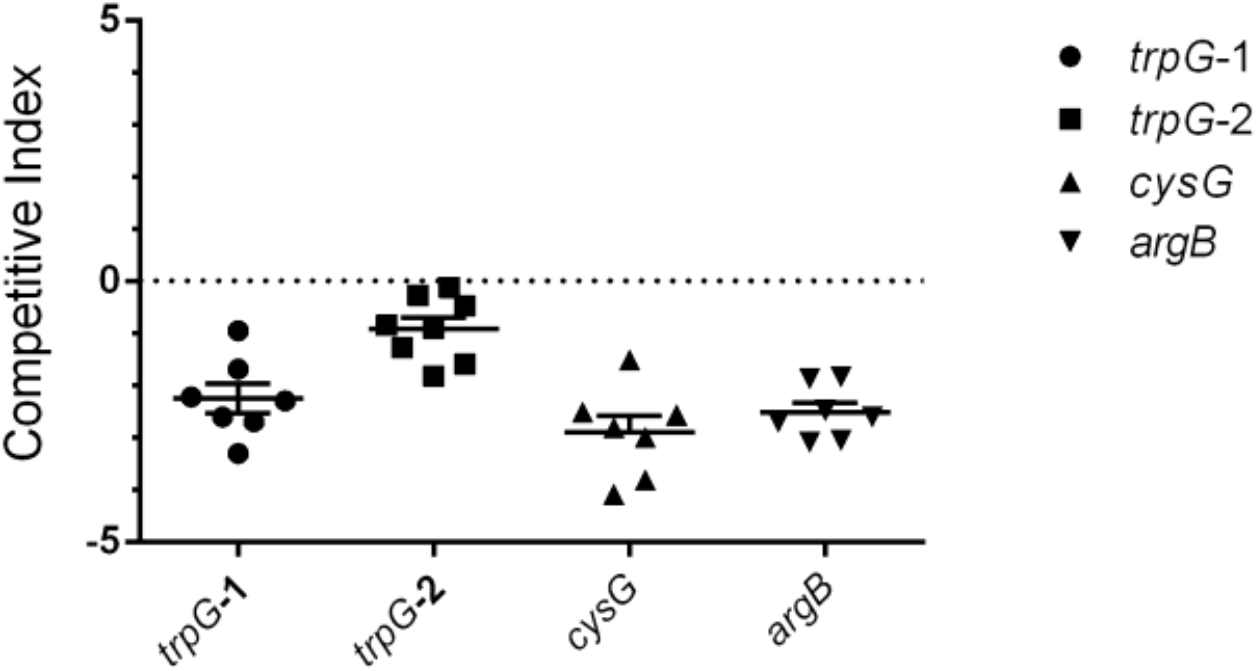
Validation of fitness determinants identified by Tn-Seq. A 1:1 ratio of wild type PAO1 and the indicated transposon mutant was administered to flies for 24 hours. Flies were then given 5% sucrose for 48 hours and homogenized at 72 hours post-challenge. Bacteria were cultured on selective and non-selective media. Each symbol represents CFU recovered from individual fly, where n= 8 flies per group and one representative replicate is shown. Competitive Index = Log_10_(CFU mutant/ CFU wild type), where a competitive index of 0 indicates equal competitive fitness.

## Discussion

Using Tn-Seq, a massively parallel transposon sequencing approach, we identified bacterial mediators of colonization for the model pathogen, *P. aeruginosa*. Functions that have been demonstrated to contribute to colonization and infection of other hosts – including flagella, exopolysaccharides, lipopolysaccharides, siderophores, and other virulence factors – were well-represented among our fly colonization candidates.

Identifying these fitness determinants yielded several insights into the lifestyle of *P. aeruginosa* in the fly. Certain colonization factors may help *P*. *aeruginosa* tolerate the stresses of the fly gut, including reactive oxygen species, low pH, and digestive peptidases. A role for oxidative stress and DNA repair in mediating bacterial survival was also indicated by a genome-wide analysis of *Franciscella novida* virulence factors in the fly (13).

Our results with regard to alginate production are surprising. We found that mutants in several pathways for polysaccharide synthesis, including alginate, were depleted in the output mixture, suggesting that alginate contributes to colonization. However, mutants in *mucA*, which encodes a negative regulator of alginate synthesis, are underrepresented in the fly, indicating that *mucA* function is important for fitness in the fly. If alginate production contributes to colonization we would have expected that the loss of the negative regulator would improve fitness. For example, MucR, a positive regulator of alginate synthesis, promotes adhesion to solid surfaces and protects *P*. *aeruginosa* from environmental stressors (www.pseudomonas.com). This unexpected result may indicate that both positive and negative regulators of alginate synthesis are required for homeostasis and optimal function. A similar relationship may explain the colonization defect in the fly associated with loss of FlgM, an anti-sigma factor that regulates flagellin synthesis in *P. aeruginosa* (www.pseudomonas.com).

The adult fly gut contains regions of low oxygen concentration, specifically an anaerobic core in the crop and a low oxygen core in part of the midgut, which is consistent with the presence of facultative anaerobes among the microbiota (37–39). As such, *P. aeruginosa* would need to adapt to an environment that is limited in oxygen in order to colonize the fly, and the loss of genes that control respiration would be detrimental. It is intriguing that in oxygen-limiting conditions, *P. aeruginosa* relies on the arginine deaminase pathway for the production of ATP (40), which may explain the requirement for arginine synthesis we observe in the fly and provide a role for amino acid synthesis during colonization that is independent of nutrition.

Amino acids and cofactors have emerged as fitness determinants in Tn-Seq studies in mice, highlighting a high degree of similarity between fly and mouse models. A prior study of *P. aeruginosa* colonization of the mouse intestine demonstrated a role for mediators of amino acid and cofactor production during infection (17). This study showed that, generally, *in vivo* gene expression and mutant fitness were not correlated; however, for genes with these functions, fitness defects were associated with increased *in vivo* gene expression, suggesting that *P. aeruginosa* requires these factors in a host environment (Table 1, [17]). In addition, purine and pyrimidine synthesis pathways have known roles in mediating exploitative competition between *P. aeruginosa* and co-colonizers of the lung and facilitating *E. coli* colonization in germ-free mice, which may explain the contribution of these factors to *P. aeruginosa* fitness (41, 42). By comparing genetic determinants of colonization in the fly to those reported for the murine intestine and in an acute burn model, we identified mediators of *P. aeruginosa* viability across host species. These functions may represent targets for therapeutic intervention (17, 34). Our results are especially notable in light of the differences in study design and statistical analysis between our investigation and these prior reports, indicating that mechanisms of *P. aeruginosa* colonization share striking elements across animal models.

Traditionally, work using *D. melanogaster* has evaluated the innate immune response to known virulence factors. However, with the advent of transposon sequencing techniques, the fly offers an opportunity for sophisticated bacterial mutant analysis in an inexpensive, genetically tractable host with a readily manipulated microbiota. Already, *Drosophila* has emerged as an important model to understand the gut microbiome and its influence on host physiology (43). These studies have revealed the diversity, composition, dynamics, and functions of microbial communities associated with flies. Recent studies have taken advantage of transposon sequencing techniques to identify microbiome factors important for colonization and impacts on the host (44–47). Of note, similar to our study, flagellar genes were identified as important colonization determinants of one microbiome member *Acetobacter fabarum* (47), suggesting this may be a common feature for bacterial persistence in the fly gut. Our previous work and other reports have highlighted a role for the microbiota in modulating enteric infection by bacteria, yeast, and viruses (48–52). We observed that colonization with a single member of the microbiota, *Lactiplantibacillus plantarum*, is sufficient to reduce mortality associated with *S. marcescens*, *P. aeruginosa* (41) and *P. entomophila* (51). Although symbiont-mediated augmentation of host defense has been proposed as the mechanism of this protective effect, a potential contribution of microbe-microbe interactions to pathogenesis has not been extensively explored in the fly (31, 53, 54). We envision subsequent Tn-Seq studies in *Drosophila* will employ fly mutants and gnotobiotic animals to interrogate the dynamic interplay among host, pathogen, and microbiota during infection.

## Materials and Methods

### Bacterial culture conditions

*P. aeruginosa* PAO1 was cultured overnight (16 h) in LB broth at 37°C with shaking at 225 rpm (50). The culture medium was supplemented with antibiotics when appropriate. *Escherichia coli* SM17-λ-pir was cultured overnight in LB with 100 µg/mL ampicillin at 37°C with shaking at 225 rpm (18).

### Fly stocks and culture

The Canton-S line of *D. melanogaster* used for these experiments was maintained on autoclaved food containing 10% dextrose, 5% heat-killed yeast, 7% cornmeal, 0.6% propionic acid, and 0.6% agar.

### Generation of *P. aeruginosa* mutant library

To perform mutagenesis, the donor and recipient strains, *E. coli* S17-λ-pir pSAM_BT20 (20) and *P. aeruginosa*, were grown separately overnight at 37°C in LB (with ampicillin (100 µg/ml) added to the *E. coli* donor). Cells from each strain were centrifuged, washed in LB, centrifuged again, and adjusted to an O.D. 600nm of 2.0. Conjugation reactions, each containing a volume of 1:3 of donor to recipient cells, were prepared on LB plates and incubated for 3 hours at 28°C. After this mating period, the conjugation reactions were resuspended in LB broth and plated on LB agar containing 50µg/mL gentamicin and 25µg/mL irgasan for 24 hours at 37°C. The next day, bacterial colonies were recovered from plates, pooled in a volume of phosphate-buffed saline (PBS) and glycerol to adjust the O.D. 600nm to 20, aliquoted, and stored at -80°C.

### Tn-Seq sample preparation, data collection, and analysis

Genomic DNA was isolated from libraries and prepared for insertional sequencing (Tn-Seq) as detailed previously (19). Samples were sequenced on an Illumina HiSeq 2000 at the Yale Center for Genome Analysis. Over 10^6^ reads per sample were obtained. Sequence analysis proceeded as described by Goodman and colleagues in (19). COG lists were accessed on May 26, 2016 from the Joint Genome Institute’s Integrated Microbial Genomes & Microbiomes dataset.

### Fly colonization for Tn-Seq analysis and characterization of mock output

To screen Tn-Seq libraries in the fly, 3-to 7-day-old Canton-S females were transferred to capillary feeders (CAFEs) (22). Flies were given access to capillaries containing 10^5^ CFU/mL of bacteria in 5% sucrose for 24 hours and capillaries containing LB and 5% sucrose for an additional 48 hours. The input consisted of the entire library without selection. Prior to culture, flies were washed with 10% household bleach, 70% ethanol, and PBS in succession. Flies were transferred to tubes with LB broth and 1.0-mm glass beads, and homogenized using a bead beater (BioSpec, Tulsa, OK). The homogenate was plated on LB agar and grown for 24 hours at 37°C. The next day, bacterial colonies were recovered from plates, pooled in a volume of LB and glycerol to adjust the O.D. 600nm to 20, aliquoted, and stored at -80°C. Each experimental replication (n=6) consisted of 250 flies. Enriched or depleted mutants were identified as previously described, using a q-value multiple hypothesis testing correction (15–18). Both depleted output conditions were independently compared against the input mutant population using an output:input abundance ratio of <1 and a significant q-value across all six replicates; enriched mutants had a output:input abundance ratio of >1 and a significant q-value across all six replicates. Prior to plating, mock output populations were maintained in capillary feeders without flies for 24 hours at the same temperature, humidity, and light levels as when the library administered to flies. The full list of genes in the library and identified in the Tn-Seq screen are listed in Tables S1-S7 of reference 21, https://figshare.com/articles/dataset/Lists_of_genes_from_PAO1_TnSeq_Assay_Miles_Manuscript/24175485.

### Fly colonization for 1:1 competition assays

Canton-S females (3-to 7-day-old) were transferred to CAFEs and given access to capillaries containing 10^5^ CFU/mL of a 1:1 suspension of wild type *P. aeruginosa* and a transposon mutant in 5% sucrose for 24 hours, then provided 5% sucrose with no bacteria for an additional 48 hours. Flies were then washed with 10% household bleach, 70% ethanol, and PBS in succession; transferred to tubes with LB broth and 1.0-mm glass beads; homogenized using a bead beater; and cultured on media with and without 10 µg/mL tetracycline. Cultures were grown overnight at 37°C for enumeration. Mutant strains (PW2175, *trpG*-1; PW2176, *trpG*-2; PW5382, *cysG*; PW9969, *argB*) were obtained from the Seattle *P. aeruginosa* PAO1 transposon mutant library.

## Acknowledgments

We thank Emily Putnam, Whitman Schofield, Andrew Goodman, Ethan Rundell, and Stephanie Shames for discussion and assistance. We thank Alexander Barron for assistance with figure design. Figures 1 and 3 were created with Biorender.com. This work was supported by the Office of the Provost at Yale University, a National Science Foundation Graduate Research Fellowship, National Institutes of Health grants 2T32GM007499-36 and P30 DK089507, and a University Grants Commission Raman Fellowship for Post Doctoral Research.

